# Genomic and transcriptomic profiles influence on brain morphology and their interactions with pain sensitivity individual differences

**DOI:** 10.1101/2024.07.30.605795

**Authors:** Yiwen Pan, Zhiguo Zhang, Xiaoke Hao, Gan Huang, Zhen Liang, Li Zhang

**Author notes:** Equal Contributions Yiwen Pan and Dr. Zhiguo Zhang. Corresponding Author Dr. Li Zhang.

## Abstract

Pain sensitivity varies widely among individuals and is influenced by a complex interplay of multi-omics factors, including genetic variations, gene expression, and brain morphology. While previous studies have identified associations between pain sensitivity and brain morphology, the exact mechanisms by which genetic profiles interact with brain structure to influence individual pain sensitivity remain unclear. In this study, we used aggregated datasets, including magnetic resonance imaging (MRI) and single nucleotide polymorphism (SNP) genotypes from 432 healthy participants, along with gene expression data from the Allen Human Brain Atlas (AHBA), to explore this multi-omics interplay. We first measured individual pain thresholds using laser stimuli and discovered structural brain differences between high and low pain sensitivity groups. We then identified two key gene sets with polarized expression patterns linked to brain morphology variations, enriched in functions related to ion channels and transmembrane transporter activities. Further statistical and mediation analyses revealed specific SNPs from *ECM1*, *SLC24A2*, and *SCN9A* genes that influence pain sensitivity, mediated through brain morphological changes in multiple basal ganglia regions. Our findings suggested that these SNPs not only affect brain structure but also modulate how individuals pain perception. Finally, we proposed an interpretation model integrating genomic, transcriptomic, and neuroimaging data, providing a detailed framework that illustrates the multi-omics contributions to individual difference in pain sensitivity. This study advances our understanding of how genetic and brain structural factors combine to shape pain perception, offering potential targets for personalized pain management strategies.

## 1. Introduction

Pain sensitivity varies significantly among people, and the individual differences can be influenced by diverse factors including genetics, cognitive states, mental health and environment.^13,16,22,65^ These factors play a crucial role in understanding underlying mechanisms of pain perception and developing effective pain management strategies. However, the complex interactions and underlying mechanisms among brain and genetic factors related to pain sensitivity remain largely unknown.

Recent research using brain imaging and genetic information has shown that alterations in brain structure^14^ and individual gene^12,55^ can lead to pain sensitivity variations. On one hand, structural magnetic resonance imaging (MRI) is a promising non-invasive modality for pain study due to its reliability.^26^ Previous MRI-based studies have shown that pain sensitivity is significantly correlated with the morphology of various brain regions, including insula, thalamus, putamen and pallidum.^14,15,23,54,60^ For example, gray matter density (GMD) have been widely used to assess the relationship between brain anatomy and pain sensitivity, revealing an inverse relationship.^14^ On the other hand, genomic studies have suggested that pain sensitivity is highly heritable, implying that it may be influenced by complex transcriptomic and genomic interactions.^18,30,43,45,67^ As an example, specific single nucleotide polymorphisms (SNPs) in genes OPRM1 have been identified to be associated with experimental pain sensitivity.^17,35^ Meanwhile, transcriptomic studies have also provided valuable insights into the expression levels of genes and their involvement in pain perception.^55^ As a bridge, imaging genetics method may reveal the link between genes and brain anatomical. These urge us to construct a model that combines brain anatomical features, transcriptomics and genomics factors for a more comprehensive investigation of pain sensitivity individual differences.

Therefore, in this work, we focus on brain imaging transcriptomics and genomics study using a self-collected dataset integrating brain imaging and genomics data, together with the Allen Human Brain Atlas (AHBA) microarray dataset.^25^ We try to explore the relationships between genetic factors and brain morphology in pain sensitivity. We aimed to address three research questions (Fig. 1): Q1. What are the transcriptomic features that are associated with pain sensitivity-related brain structural alternation? Q2. What are the brain phenotype differences among various SNP genotyping and pain sensitivity groups? and Q3. What are the possible interactions among genetic factors, brain factors and pain sensitivity?

**Fig. 1.**
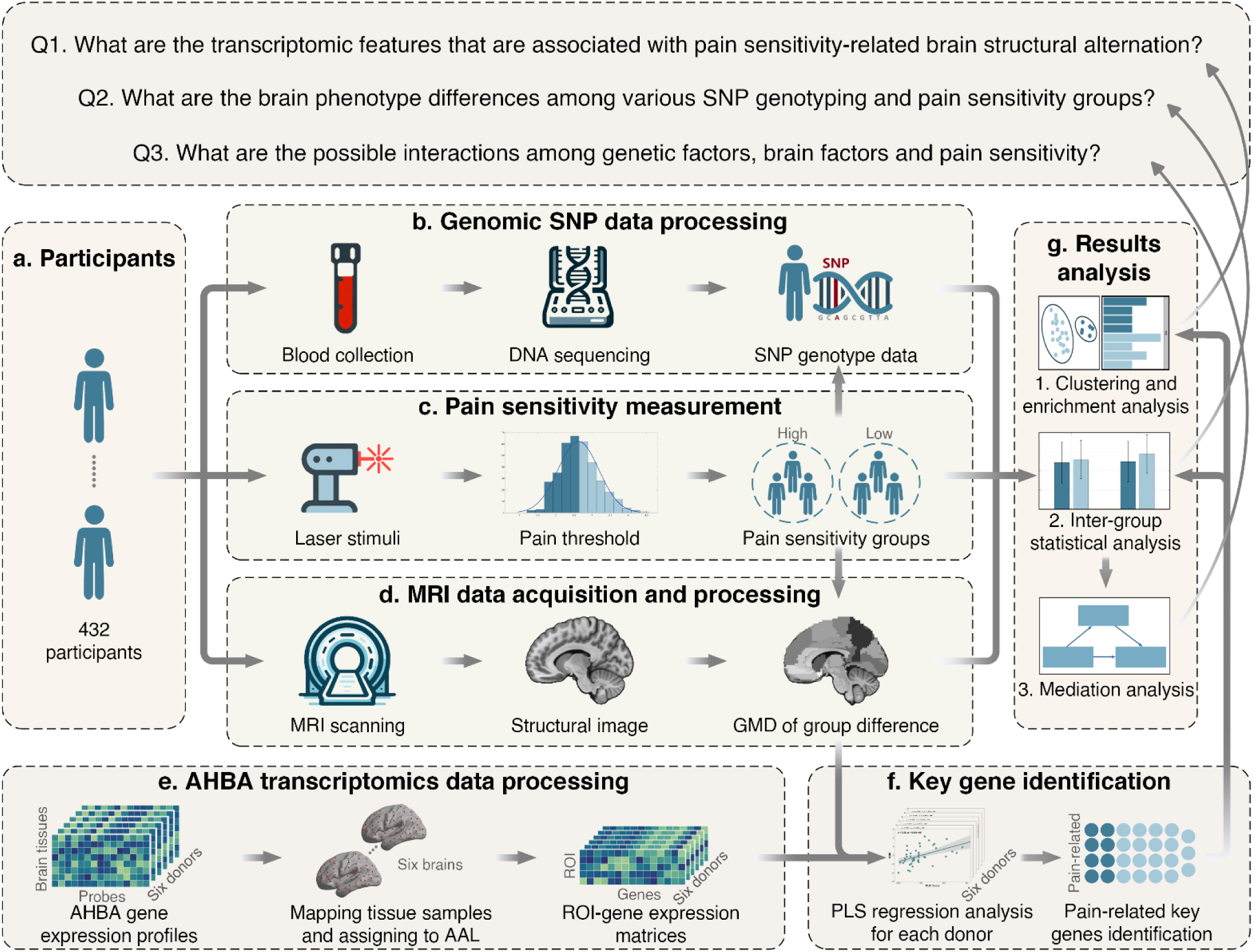
The research questions and the overall flowchart of this study. This study aims to answer three questions (Q1 to Q3). The data collection and analysis methods are as follows: (a) First, in total 432 healthy participants were enrolled in this study. (b) Blood samples were collected to obtain the SNP genotype data. (c) The individual pain sensitivity was measured via laser stimuli and the subjects were partitioned into high and low sensitivity groups. (d) Brain structural images were obtained by MRI scanning and the GMD differences between high and low sensitivity difference were calculated. (e) AHBA gene expression data were processed to get ROI-gene expression matrices for six donors. (f) By using partial least squares regression to integrate brain imaging GMD group difference features and gene expression features, pain-related key genes were identified. (g) 1. Clustering and enrichment analysis were conducted for the key genes to answer question Q1; 2. By incorporating the SNP genotype of the key genes, statistical analysis was applied to reveal inter-group differences in terms of pain sensitivity and genotypes to answer question Q2; 3. and finally mediation analysis was applied to discover possible relationships among gene factor, brain factor and pain sensitivity to answer question Q3.

To answer these questions, in total of 432 healthy participants were enrolled and their pain sensitivity were measured using laser stimuli. MRI and SNP data were gathered to examine the brain imaging and genomics patterns. The inclusion of the AHBA microarray dataset provides valuable insights into how alterations in gene expression are associated with macroscale brain phenotypes (brain imaging) and microscale genotypes (SNP). We employed comprehensive methods on multi-omics datasets to identify key gene sets that are associated with pain sensitivity. Further, we tried to uncover the mediation pattern between microscale pain sensitivity-related SNP and macroscale brain anatomical features. Finally, we integrated the genomics, transcriptomics and neuroimaging phenotypes to build an interpretation model, which has the potential to reveal candidate genes that contribute to pain sensitivity individual differences.

## 2. Materials

### 2.1. Participants

A total of 432 healthy participants (259 females, average age: 21.3 ± 3.99 years) were recruited for this experiment via college and community advertisements. Twenty left-handed participants were excluded from the analysis (412 remained) due to differences in brain structure and function observed between left-handed and right-handed individuals.^39,48,64^ Before the experiments, careful screening procedures were conducted to ensure that participants had no history of chronic pain, neurological conditions, cerebrovascular diseases, coronary heart disease, or mental health disorders. Additionally, participants were verified to have no contraindications for undergoing MRI examinations. The study obtained approval by the Ethics Committee of the Institute of Psychology, Chinese Academy of Sciences. All participants provided written informed consent before their participation. The data collected in this study are not available to the public, as the participants did not consent to the public dissemination of their information. An outline illustrating the data collection and analysis methods utilized in the study is summarized in Fig. 1.

### 2.2. Pain sensitivity definition

We used the pain threshold to define one’s pain sensitivity. Unlike other criteria such as pain tolerance, pain threshold is more stable within an individual.^57,68^ Specifically, this study rated the participants’ pain sensitivity by using a laser pain threshold before the MRI scan. We first recorded the skin temperature of each participant. A series of infrared neodymium:yttrium-aluminum-perovskite (Nd:YAP) stimuli were applied to the area between the thumb and index finger on the back of the participant’s left hand. To prevent skin receptor fatigue due to repeated laser stimulation, the laser position was randomly changed after each stimulus. The process began with an energy level of 1 J, incrementing by 0.25 J for subsequent stimuli. After each stimulus, participants rated their pain experience on a scale ranging from 0 (no pain at all) to 10 (unbearable pain). When a participant reported a rating of 4, the corresponding energy level was recorded as their laser pain threshold. Each participant underwent two separate measurements within a 1-hour interval, and the laser pain threshold was calculated by averaging these measurements. The laser pain threshold values underwent z-score normalization to establish the pain sensitivity threshold to meet a normal distribution. To make the high and low sensitivity groups more distinguishable, we used the median value of the laser pain threshold as the primary ranking, and skin temperature as the secondary ranking (since it has been proven to be related to pain sensitivity^47,66^) as a benchmark and excluded 10% of total participants around this benchmark from the analysis. The remaining individuals’ skin temperature was ranked in descending order with their pain thresholds were sorted in ascending order from low pain thresholds (high sensitivity) to high pain thresholds (low sensitivity). Based on this sorted order, the participants were divided into high/low pain sensitivity groups.

### 2.3. MRI data acquisition and preprocessing

MRI data were acquired using a 3.0-Telsa system (GE Medical Systems, Milwaukee, WI, United States) at the Brain and Cognitive Neuroscience Research Center, Liaoning Normal University, Dalian, China. A standard 8-channel head coil and restraining foam pads was used. Structural T1-weighted images were obtained using a 3D magnetization-prepared rapid gradient echo sequence with the following imaging parameters: flip angle = 8◦, field of view = 256 × 256 mm2, data matrix = 256 × 256, in-plane resolution = 1 × 1 mm2, slices = 176, and slice thickness = 1 mm. During the scan, participants were instructed to remain awake, and not think about anything with their eyes closed.

Preprocessing steps were carried out utilizing CAT12 toolboxes with default settings within SPM12 (Statistical Parametric Mapping). Voxel-based morphometry (VBM) methodology^21^ was employed to process and extract brain-wide MRI imaging phenotypes of interest. The structural images underwent segmentation into gray matter (GM), white matter (WM), and cerebrospinal fluid (CSF) components by applying registration to the Montreal Neurological Institute (MNI) stereotactic space, followed by nonlinear deformation. The calculation of nonlinear deformation parameters was achieved using the built-in high-dimensional Diffeomorphic Anatomical Registration Through Exponentiated Lie Algebra (DARTEL) algorithm.^2^ The resultant warping functions generated by DARTEL were employed to spatially normalize the GM segments, which were then modulated by the Jacobian determinant. Subsequently, the normalized GM images underwent smoothing using an 8-mm full width at half maximum (FWHM) Gaussian kernel. After preprocessing the MRI data, the smoothed GM image was obtained. The automated anatomical labeling (AAL) atlas^62^ was used to extract the GMD from different regions of interest (ROIs) for each participant.

### 2.4. Genomic SNP data

Five-milliliter blood samples were obtained from participants for DNA extraction. Genotypes of SNPs were determined using the CapitalBio Technology Corporation Precision Medicine Research Array based on the Affymetrix platform. A total of 748,659 SNP markers were collected as genome-wide genotype information. To handle possible defects of genetic data, the following quality control (QC) steps were performed using the PLINK software package (v1.9).^50^ Firstly, only SNPs located on autosomes were included, and SNPs on the sex chromosomes (27,932 SNPs) were removed. Secondly, SNPs were excluded if they could not meet any of the following criteria: (1) call rate per SNP ≥ 90% (46,803 SNPs), (2) minor allele frequency (MAF) ≥ 5% (319,751 SNPs), and (3) Hardy-Weinberg equilibrium test of p ≤ 10e-6 (752 SNPs). Participants were excluded from the analysis if any of the following criteria was not satisfied: (1) call rate per participant ≥ 90%; (2) gender check (5 participants were excluded, 407 remained). Population stratification analysis^19^ was performed by applying the principal component analysis (PCA), which is considered to be a very useful method of adjusting spurious associations. This method allows for the extraction of ethnic information from the sample and the identification of potential ethnic outliers. After the QC procedure, 407 participants and 353,421 out of 748,659 SNP markers were retained in the analysis.

### 2.5. Transcriptomic gene expression data

The microarray-based gene expression data used in this study were sourced from AHBA (http://human.brain-map.org).^25^ This dataset contains expression data derived from six human postmortem donors, comprising one female and five males aged between 24 and 57 years. More precisely, the AHBA dataset contains the expression of 20,737 genes from the entire brain, with each gene represented by unique Entrez IDs measured through 58,692 probes across 3,702 spatially different tissue samples. The *abagen* toolbox^40^ was used to process the microarray expression data, which offers standardized workflows for the AHBA dataset. The gene expression data underwent preprocessing steps as detailed in a previous study^1^: (1) Microarray probe-to-gene mappings were reannotated using the latest National Center for Biotechnology Information (NCBI) database, with probes that didn’t match a valid Entrez ID discarded. (2) Probes with expression intensities lower than the background noise in over 50% of samples were filtered out. (3) In cases where multiple probes indexed the expression of the same gene, a representative probe exhibiting the highest pooled correlation across donors was selected. (4) Matching of microarray samples to brain regions was achieved using MNI coordinates, considering coordinates within 2 mm of the region boundary. (5) Gene expression values were normalized within each donor to address the inter-individual differences. This normalization involved applying the scaled robust sigmoid function and rescaling to the unit interval. (6) Genes were further refined based on differential stability criteria to retain only stable genes for subsequent analysis.^24^

It’s important to note that the samples from each donor were averaged separately rather than across all six donors. Additionally, because only two donors out of six comprise brain samples from the right hemisphere, we only used the left hemisphere for further analysis.^1^ Following these processing steps, the expression values became more comparable across donors, resulting in the acquisition of six gene expression matrices at different donor levels for subsequent analysis of connecting neuroimaging phenotype.

## 3. Methods

### 3.1. Brain imaging phenotype extraction

In our study, we first conducted a comparison of brain imaging phenotype characteristics between individuals with high and low pain sensitivity groups. Differences in GMD features among various ROIs were calculated using two-tailed two-sample t-tests with a significance level of p < 0.05, corrected by the Benjamini-Hochberg false discovery rate (BH-FDR) method. We also computed the average phenotype values to get a phenotype representation vector for each group. Afterward, we performed a vector subtraction, yielding a differential phenotype vector between high and low pain sensitivity groups for further regression analyses.

### 3.2. Imaging transcriptomics analysis

A partial least squares (PLS) regression was employed to explore the relationship between the gene expression patterns of each donor and the brain phenotype. PLS regression is capable of predicting multiple response variables (brain phenotype values) from multiple predictor variables (gene expression data) by establishing an interrelation between them.^28,34,41^ In our study, we first obtained the brain expression level of each gene from each ROI in each donor using preprocessed AHBA data. Next, both the gene expression levels and GMD difference values were normalized by subtracting the mean and dividing by the standard deviation (SD) across all samples. To evaluate the significance of GMD features in the model, we computed Variable Importance in Projection (VIP) scores. VIP scores represent a measure of the importance of variables in multivariate data analysis, indicating the degree to which a variable contributes to explaining variations in the response variables.^8^ Variables with VIP scores greater than 1 were considered significant in predicting the PLS regression model and were extracted from the previously mentioned gene expression matrix. As a result, this process led to the acquisition of six matrices of transcriptional levels (45 regions of 6 donors multiplied by corresponding gene expression levels) and a set of differential GMD values (45 regions multiplied by brain phenotypes).

In our PLS regression analysis, the predictor variables comprised normalized matrices representing average gene expression levels across all regions for each donor level (45 × 5333; 45 × 5439; 45 × 5089; 45 × 5840; 45 × 5700; 45 × 5286). Meanwhile, the response variables were the normalized vectors of average differential GMD phenotype across all regions (45 × 1). The output of components from PLS regression was ranked based on the explained variances between the predictor and response variables, with particular emphasis on the first PLS component (PLS1). PLS1 aimed to provide an optimal low-dimensional representation of the covariance between two high-dimensional datasets, namely gene expression and differential phenotype values.^34^

To examine whether the explained variance of PLS1 significantly exceeded chance expectations, we conducted permutation tests (10,000 times).^31,44^ This involved multiple iterations of shuffling data labels and performing recalculations of the PLS regression analysis to establish a null distribution of explained variances. Furthermore, we used bootstrapping techniques to estimate the significance of individual genes contributing to the PLS components. Specifically, after obtaining PLS1 weights for each gene, we performed 10,000 resampling iterations with the replacement of 45 brain regions to re-conduct the PLS regression analysis at the donor level via bootstrap tests. This process generated weights for each gene. The ratio of each gene’s weight to its bootstrap standard error was computed to derive z-scores (PLS regression weights divided by their standard errors). These z-scores were transformed into p-values to evaluate the contribution of each gene to PLS1. To identify significant genes, we applied the BH-FDR correction procedure, setting a significance threshold at p-value < 0.05.

To address potential biases arising from transcriptomic variations among the six donors, we constructed separate PLS regression models for each donor. Through this approach, we aimed to eliminate donor-specific influences on the analysis. Specifically, following the creation of separate PLS regression models for each donor, we identified genes that demonstrated noteworthy associations with differential phenotype values linked to pain sensitivity. Subsequently, these genes that are consistently found across all donors were selected for subsequent investigation and further analysis.

### 3.3. Key gene identification

By comparing the significant genes identified across all six donor levels, we identified key genes strongly associated with GMD variations related to pain sensitivity. These key genes were found to have functional information related to pain sensitivity based on the data processing steps employed. Subsequently, we utilized the gene expression features of these key genes across 45 brain regions at six donor levels for conducting *k*-means clustering. To determine the quality and coherence of the resulting clusters, the silhouette measurement was used to get the optimal classification groups by calculating the relative intra-cluster similarity and inter-cluster dissimilarity.^52^ We further calculated the correlation between each gene based on its expression value of all ROIs and all donors from the AHBA dataset. Thus, the expression value vector was obtained by concatenating 45 ROIs’ expression value of all AHBA donors, resulting in a 45 × 6 = 270 long vector. Pearson correlation coefficients were then calculated based on the expression value vector to determine the relationships among all genes.

Furthermore, we explored potential functional differences between the identified groups using the Gene Ontology (GO) database.^3^ Specifically, we conducted GO analyses to comprehensively examine the functional roles of the identified genes concerning molecular function, biological processes, and cellular components. These GO analyses of the key genes were executed utilizing the *clusterProfiler* package within the R software, employing a significance threshold set at a p-value of 0.05.

We established a comprehensive pain gene set by integrating pain-related genes sourced from multiple databases, including the Human Pain Genetics Database,^42^ Pain Genes Database,^29^ Gene Ontology search employing the keyword “pain”, and Pain Research Forum from https://www.iasp-pain.org/publications/pain-research-forum/. We compared the identified key genes related to GMD from our analysis with these established pain gene sets to highlight significant pain-related genes associated with GMD variations. We then extracted the expression values of these genes across 45 ROIs to examine the gene expression pattern at the donor level.

### 3.4. SNP genotyping analysis

By integrating the self-collected preprocessed genotype SNP data, we selected target SNPs corresponding to the identified genes from previous step. Additional statistical analyses were performed aimed at quantitatively determining the differences in phenotype among various genotype groups. To handle the uneven distribution of participants across different groups, one of the homozygous groups with a smaller participant count was merged with the heterozygous group, resulting in the formation of a risk allele-carrier group.^33^ Meanwhile, the participants were divided into high and low pain sensitivity groups, enabling us to conduct a two-way ANOVA (2 × 2 ANOVA: pain sensitivity × genotypes). It was executed to evaluate the combined impact of the target SNPs and pain sensitivity on GMD differences within the significant ROIs. All statistical analyses were performed using SPSS 22.0 software.

### 3.5. Mediation analysis

To further investigate the influence of GMD on the relationship between genetic factors and pain sensitivity, we performed a mediation analysis to determine if the association between variables (SNP and pain sensitivity) can be partially explained by the effect of a mediating variable (GMD).^37^ In this study, the genotype of each SNP served as the independent variable, pain sensitivity measurements as the dependent variable, and differential GMD values of significant ROIs as the mediating variable. The hypothetical mediation models were conducted using path analysis in structural equation modeling (SEM) with maximum likelihood estimation. In addition, a bootstrapping sampling procedure was employed to calculate bias-corrected 95% confidence intervals (CIs) for evaluating the significance of indirect and direct effects in the mediation model.^38^ Bootstrapping is a nonparametric resampling procedure, and as such, it does not violate assumptions of normality. In this study, statistical significance was considered when the 95% CI (based on 5,000 bootstrap samples) excluded 0 between the lower and upper limits.^49^ Standardized estimate (β), standard error (SE), 95% CI, and p-value were reported for both direct and indirect effects. All mediation analyses were performed with Amos (Version 24.0, IBM Corp, Armonk, NY).

## 4. Results

### 4.1. Participants characteristics

After excluding 10% subjects using pain threshold values and skin temperatures as the benchmarks, 366 out of the remained 407 participants were kept in the present study. These pain thresholds were arranged from low (high pain sensitivity) to high (low pain sensitivity). The participants were categorized into 2 groups: 157 individuals in the high pain sensitivity group and 209 individuals in the low pain sensitivity group. Subsequently, the pain thresholds underwent normalization via z-score transformation. Table 1 provides an overview of the demographic details of the sample analyzed in this study. A noteworthy observation from the independent-sample t-test regarding structural phenotypes suggests a significant difference (p-value = 0.003) in gender distribution among the participants categorized into high and low sensitivity groups. In our following analysis, the sex factor is regressed as a covariate aiming to control for its potential influence on the observed results.

**Table 1.** Demographic information and the total number of participants are included in each analysis.

| Category | High | Low | p-value |
| --- | --- | --- | --- |
| Number of participants | 157 | 209 | - |
| Gender (M/F)<sup>a</sup> | 48/109 | 96/113 | 0.003 |
| Age (years, mean ± SD)<sup>c</sup> | 21.3 ± 4.4 | 21.4 ± 4.0 | 0.899 |
| Education (years, mean ± SD)<sup>c</sup> | 14.6 ± 2.1 | 14.5 ± 1.9 | 0.633 |

<sup>a</sup>Chi-square test. <sup>b</sup>p-value <0.05. <sup>c</sup>Two-sample t-test.

### 4.2. GMD associated with pain sensitivity

We compared the GMD differences in 45 brain regions between the high and low pain sensitivity groups. As a result, we identified 15 ROIs that exhibited significant inter-group differences, with the statistical significance corrected using the BH-FDR method. The results are shown in Fig. 2. Among these 15 ROIs, we found many of them belong to the pain matrix or are highly related to pain perception. These ROIs include insula (p-value = 0.014), amygdala (p-value = 0.020), putamen (p-value = 0.014), pallidum (p-value = 0.026), and thalamus (p-value = 0.021). This particular discovery suggests that individuals in the low pain sensitivity group have higher GMD compared to those in the high pain sensitivity group.

**Fig. 2.**
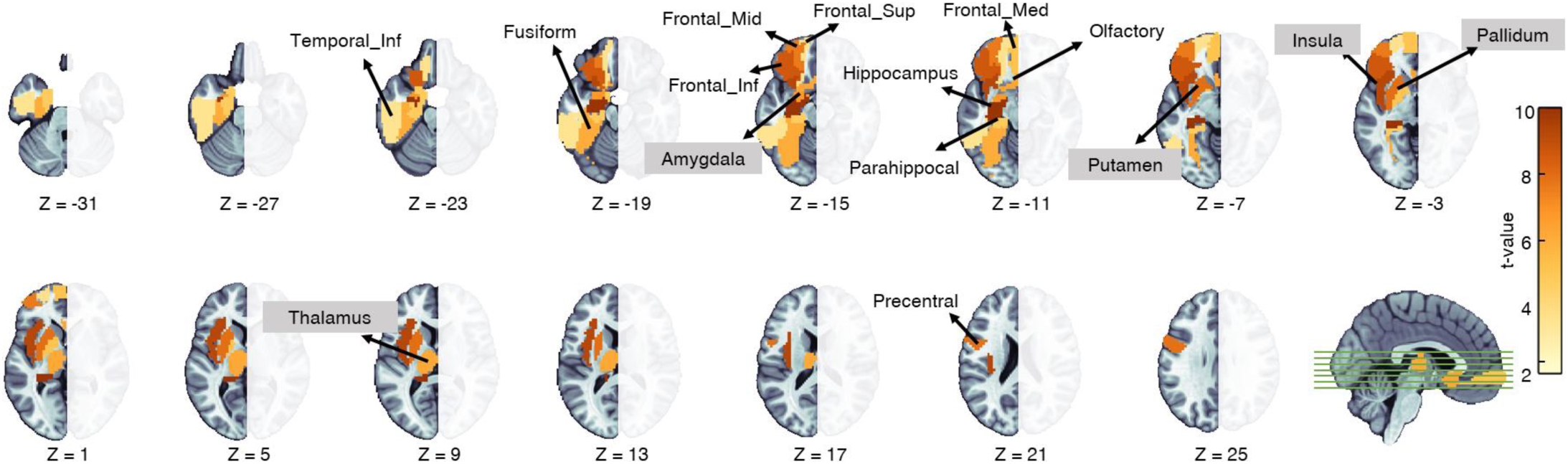
Identified 15 ROIs that exhibited significant differences between high and low pain sensitivity groups in terms of GMD. The ROIs with gray backgrounds were commonly known to be belonging to the “pain matrix”. Since only the left hemisphere of the AHBA dataset was considered in this study, here we only show identified ROIs of the left brain.

### 4.3. Correlations between gene expression and GMD

PLS regression were used to identify patterns of gene expression that exhibited correlations with the GMD differences across six donors. The results are presented in Fig. 3. Firstly, Fig. 3(a) illustrates the correlations between the PLS1 scores and the GMD difference values. The scatter plots demonstrate significant positive correlations between the PLS1 scores and the GMD values for each donor. The Pearson’s correlation coefficients ranged from 0.49 to 0.63, demonstrating a significant relationship between these variables (all p-values < 0.001). These results imply the role played by PLS1 in capturing the association between gene expression patterns and GMD differences related to pain sensitivity across different donors. Secondly, as shown in Fig. 3(b), the permutation tests revealed that the PLS1 scores explained a significant amount of the variance in GMD across all six donors. The genes from each donor were repeatedly identified to make substantial contributions to PLS1 (permutation test, p-value < 0.05), significantly more than expected by chance. Notably, the PLS1 component independently accounted for the highest observed variance, explaining a minimum of 38.8% of the total variance.

**Fig. 3.**
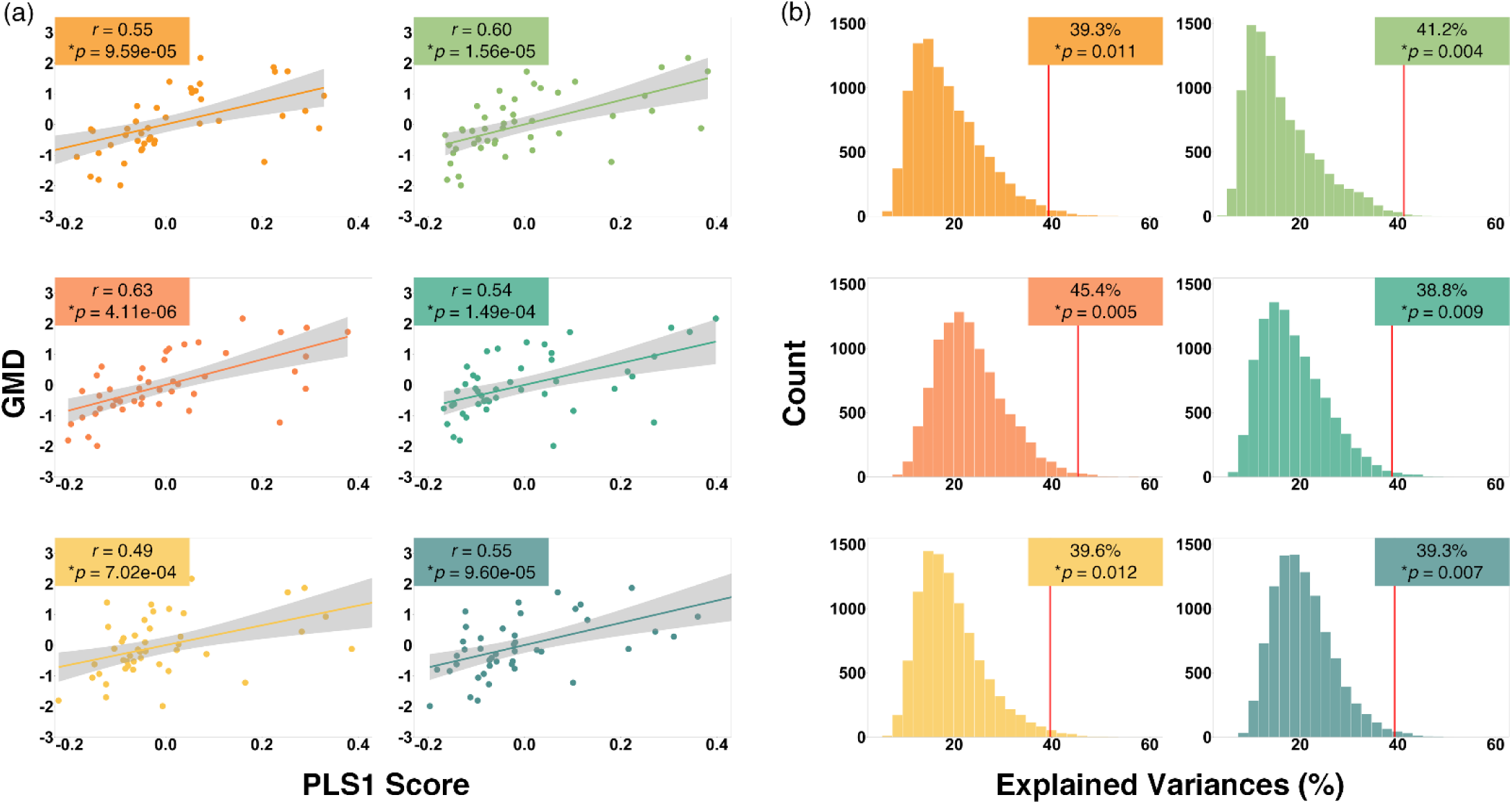
(a) The results from PLS regression analysis for 6 donors. The correlations between PLS1 scores and the GMD differences values are shown in each subfigure. (b) The results from permutation tests for 6 donors.

### 4.4. Key gene clustering and enrichment analysis results

Furthermore, we identified genes that significantly contributed to PLS1 consistently present across all six donors using the bootstrapping method. Specifically, we identified a gene set that is associated with pain sensitivity-related GMD alternation for each donor. In order to obtain a relatively stable result, we only selected genes that were identified from all six donors for further analysis. In the end, 27 key genes were identified to be significantly associated with differences in GMD between two pain sensitivity groups and were used. The gene identification and overlapping situations are demonstrated as a Venn diagram in the supplementary materials, together with 27 key gene names and other related information.

Next, we used a *k*-means clustering method on these 27 key genes and the results are shown in Fig. 4(a). To compare the clustering effect with different *k*, the cluster numbers were chosen as 2 and 3. The silhouette coefficients are 0.713 and 0.531 for *k* = 2 and 3 respectively. These results indicated that the optimal division for these 27 genes lies in categorizing them into two distinct clusters instead of three. Following the clustering analysis, the 27 significant genes were divided into two groups: group 1 with 21 genes, and group 2 with 6 genes. Further, the gene expression correlation results are also shown in Fig. 4(b) where red indicates positive correlations and blue indicates negative correlations (corrected by the BH-FDR method, p-value < 0.001).

**Fig. 4.**
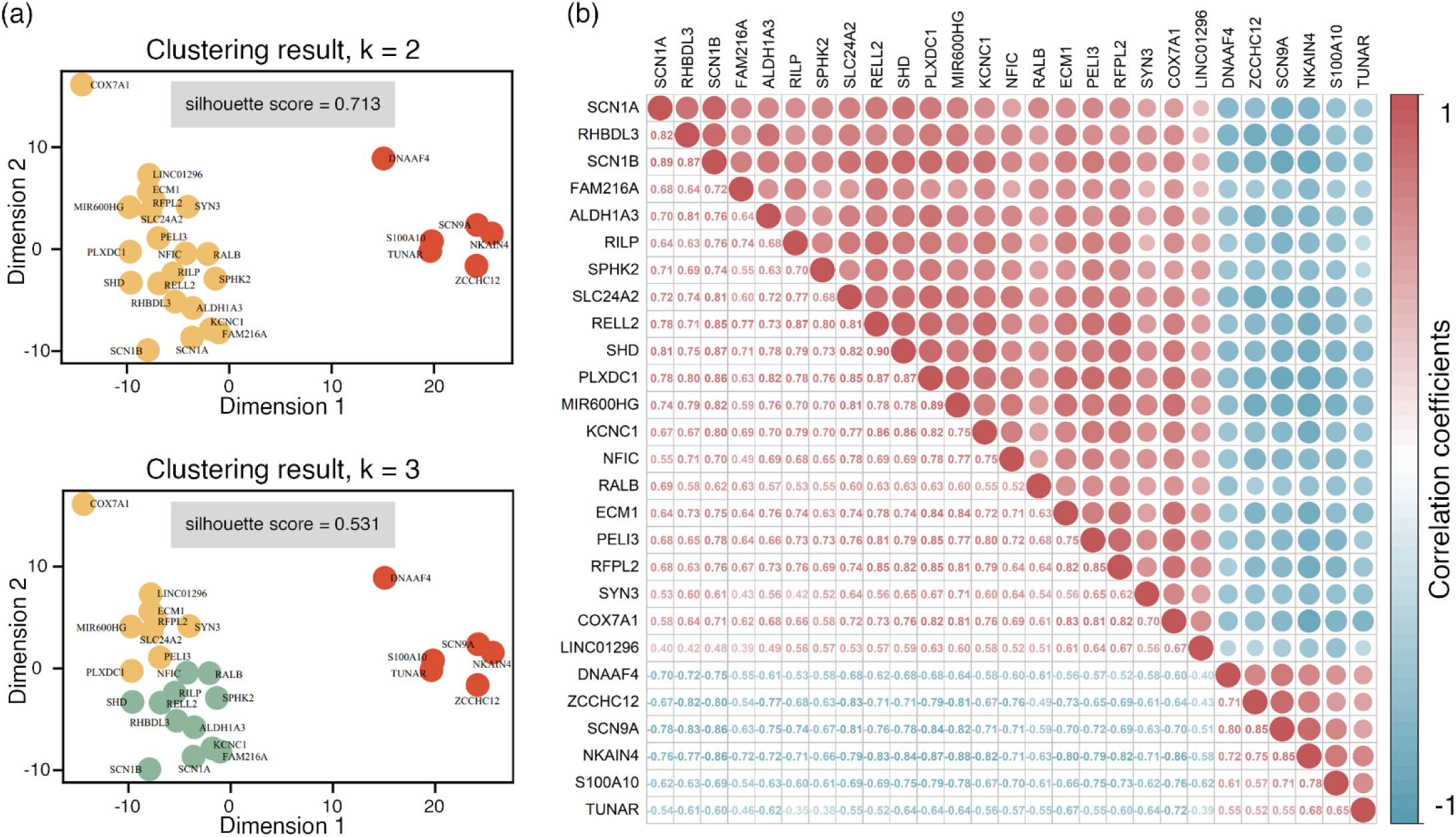
(a) The *k*-means clustering results for 27 key genes by choosing *k* = 2 (top) and *k* =3 (bottom) respectively. (b) The correlation coefficient matrix among 27 identified key genes.

Figure 5(a) demonstrates the gene groups and gene network in which only high correlation (absolute value greater or equal to 0.6) connections were kept. Next, the GO enrichment analysis was conducted for each group to examine the functional characteristics associated with these genes. The results are shown in Figs. 5(b) and (c) (all p-values < 0.05). First, Fig. 5(b) shows the GO enrichment analysis for the 21 genes within group 1. These genes exhibit significant over-expression in terms of molecular functions and cellular components. The significantly enriched molecular functions contain monovalent inorganic cation transmembrane transporter activity, voltage-gated sodium channels (VGSC) activity, cation channel activity, sodium and metal ion transmembrane transporter activity and ion and sodium channel activity. Meanwhile, in terms of cellular components, the enrichment predominantly occurs in areas such as the main axon, VGSC complex, node of Ranvier, sodium and cation channel complex. On the other hand, Fig. 5(c) shows the GO enrichment analysis outcomes for the 6 genes within group 2. These genes can only be enriched in cellular components. Specifically, these genes are enriched in the VGSC complex and sodium channel complex.

**Fig. 5.**
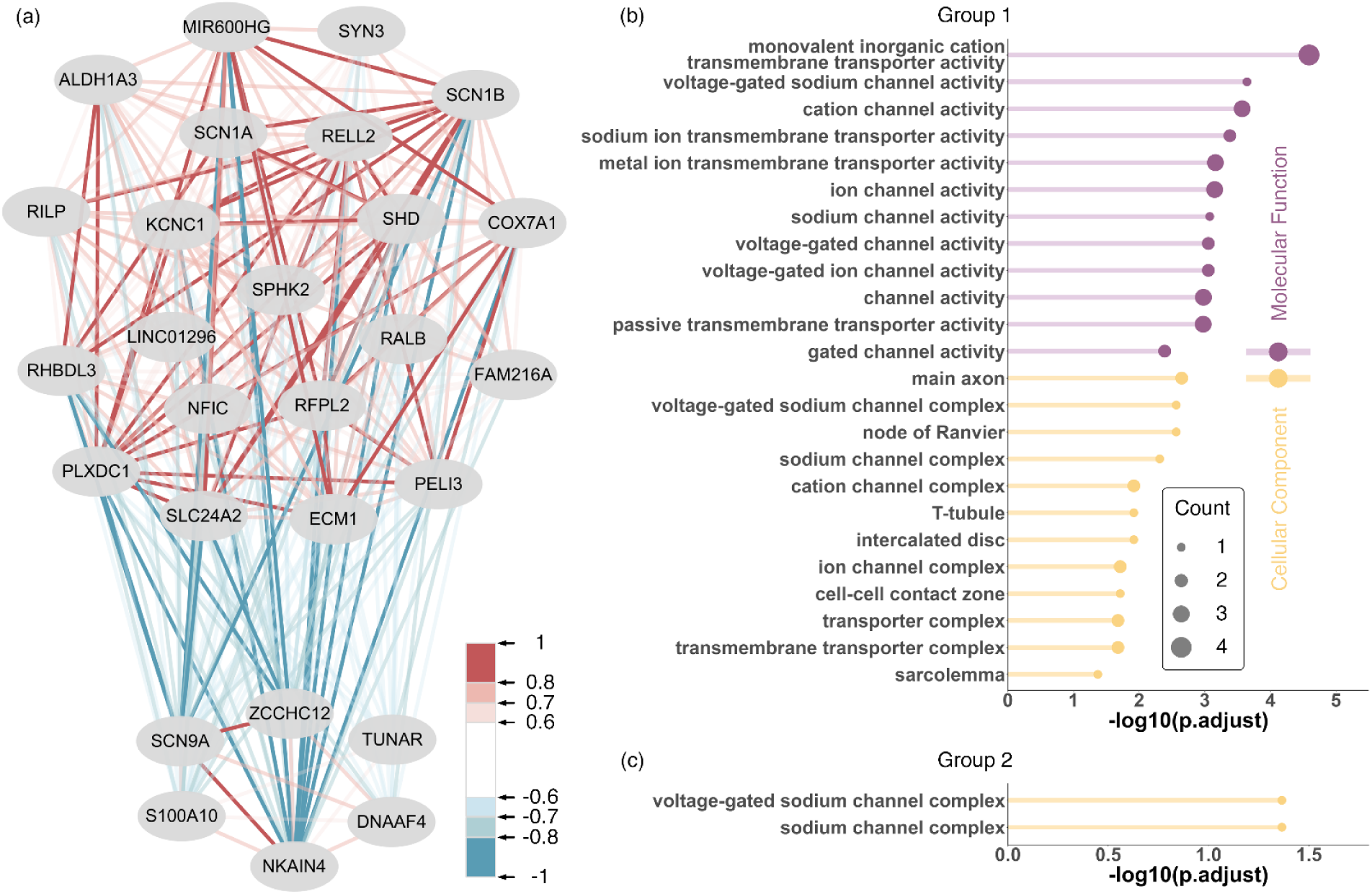
(a) The clustering results for 27 key genes. These genes can be clustered into two groups. The upper group is group 1 with 21 genes and the lower group is group 2 with 6 genes. The lines between each gene pair indicate the expression correlation among all ROIs, where red indicates a positive correlation and blue indicates a negative correlation. The correlation value is indicated as the color bar. (b) and (c) The GO analysis results for genes in groups 1 and 2 respectively.

### 4.5. Pain-related key gene expression

By taking the intersection set of these 27 key genes and the aforementioned pain-related genes set from multiple database sources (including the Human Pain Genetics Database,^42^ Pain Genes Database,^29^ Gene Ontology search employing the keyword “pain”, and Pain Research from https://www.iasp-pain.org/publications/pain-research-forum/), we further narrow our analysis down to 8 significant pain-related genes: *ECM1*, *SCN1A*, *SCN1B*, *SLC24A2*, *SPHK2*, *SYN3*, *S100A10* and *SCN9A*. More details of this intersection can be found in the supplementary document. Fig. 6 shows the expression values of these 8 genes on 45 ROIs. In the figure, the circle size is determined by the expression level among all genes and all ROIs. For each row (gene), the expression is respectively ranked from largest to smallest. The ranking order is indicated below each circle. The color of each circle represents the brain region in the AAL atlas of the corresponding ROI (blue: central region, green: frontal lobe, orange: insula, cyan: limbic lobe, red: occipital lobe, gray: parietal lobe, purple: subcortical nuclei, and yellow: temporal lobe). The ROIs with a gray background are brain regions that belong to the “pain matrix”, including the dorsolateral prefrontal cortex (dlPFC), insula, anterior cingulate cortex (ACC), S1, S2 and regions from basal ganglia (amygdala, caudate, putamen, pallidum and thalamus). Based on the previous *k*-means clustering result, the first 6 genes belong to group 1 (darker colors) and the other 2 genes belong to group 2 (lighter colors). The bar plot in Fig. 6 indicates the normalized gene expression of these 8 genes at different brain regions. The overall bar height is the total expression level of all 8 genes. Within each bar, the darker color represents the expression ratio of the first 6 genes (group 1) and the lighter color represents the other 2 genes (group 2).

**Fig. 6.**
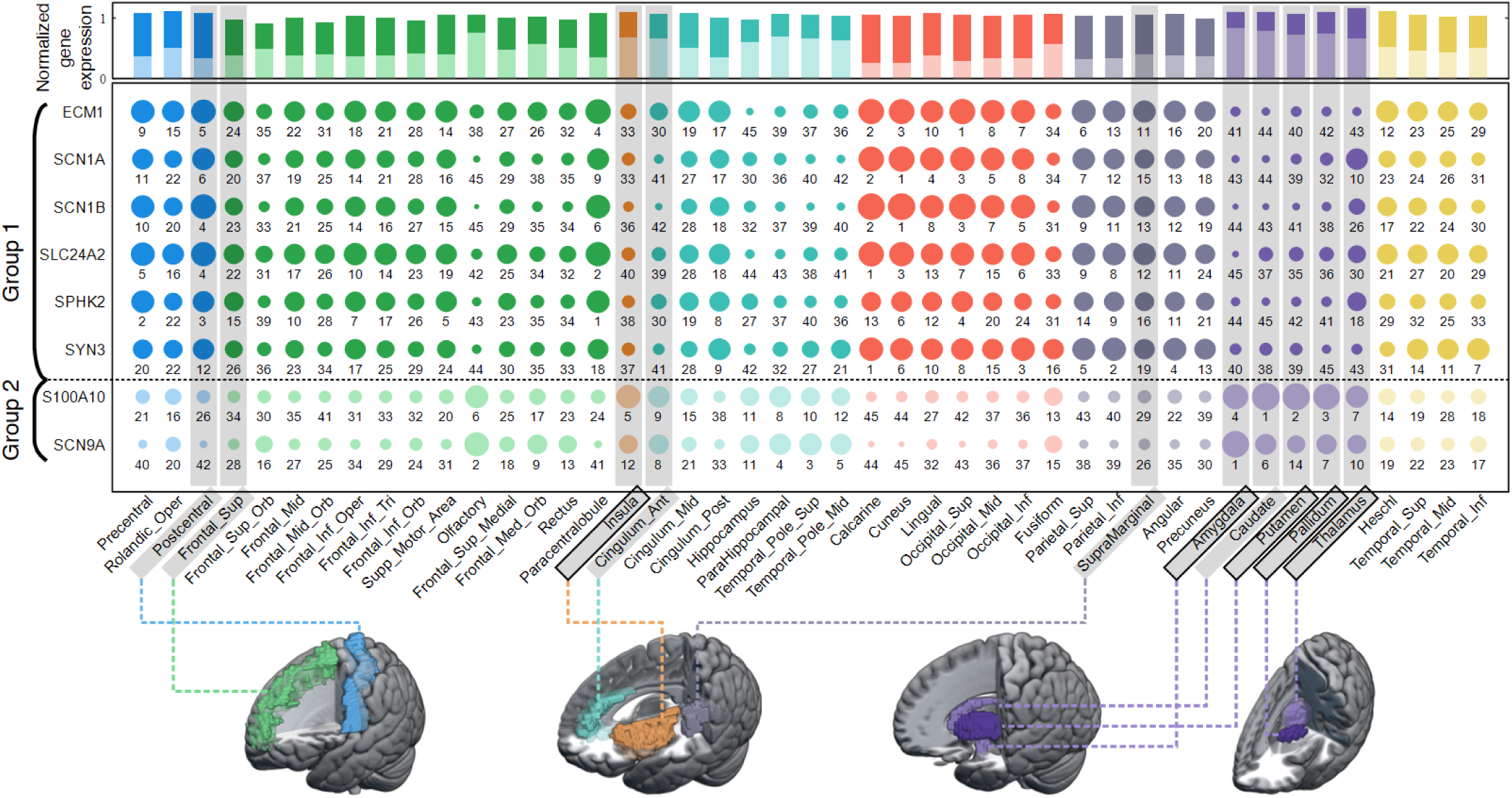
The expression values of 8 key genes (*ECM1*, *SCN1A*, *SCN1B*, *SLC24A2*, *SPHK2*, *SYN3*, *S100A10* and *SCN9A*) at different brain regions. The circle size is determined by the expression level among all genes and all ROIs. For each row, the expression value is ranked from largest to smallest and the ranking order is listed below the circle. The bar plot indicates the normalized gene expression of these 8 genes at different brain regions, where the darker part is from group 1 (genes 1 to 6) and the lighter part is from group 2 (genes 7 and 8). Brain regions with gray backgrounds are known to belong to the “pain matrix”.

From this figure, it is interesting to observe that the regions from basal ganglia (shown in purple) show a polarized expression pattern. Specifically, genes in group 1 (*ECM1*, *SCN1A*, *SCN1B*, *SLC24A2*, *SPHK2* and *SYN3*) have small expression values, and genes in group 2 (*S100A10* and *SCN9A*) have large expression values in these areas. It is especially true for amygdala, caudate, putamen and pallidum. By comparing the gene expression level with other brain regions, genes in group 1 from basal ganglia have almost the smallest expression values among all 45 ROIs, and genes in group 2 have the largest expression value. For example, the top 4 out of 5 top-expressed ROIs of gene S100A10 are in the basal ganglia region. On the other hand, these 4 ROIs have the smallest expression value for the SPHK2 gene. To verify the clustering result, we also conducted the *k*-means clustering method based on the 8 identified key genes based on whole-brain 45 ROIs and pain matrix ROIs (shown in gray background in Fig. 6). Interested readers may refer to the supplementary document for more information.

### 4.6. SNP genotyping statistical results

Further, by integrating our self-collected genotyping dataset, we investigated SNPs of the aforementioned 8 pain-related key genes, revealing 3, 21, 0, 51, 0, 86, 3 and 27 SNPs for them respectively. Subsequently, we conducted a two-way ANOVA analysis on all these SNPs to examine the relationships between the genotypes and pain sensitivity on GMD in significant brain regions. The analysis contains 15 previously mentioned important ROIs that exhibited significant differences in GMD between high and low pain sensitivity groups, including the insula, hippocampus, fusiform, amygdala, putamen, pallidum, thalamus, and so on. As a result, we identified SNPs from 3 genes, namely *ECM1*, *SLC24A2* and *SCN9A* with significant statistical results.

For the *ECM1* gene, there are 3 SNPs included in our dataset. After performing the statistical analysis, one SNP rs13294 showed significant results between pain sensitivity and genotyping groups. The results are shown in Figs. 7(a) and (b). There are in total 366 participants (156 with high pain sensitivity: 78 AX, 78 GG; 208 with low pain sensitivity: 86 AX, 122 GG; 2 missing values). Due to the limited number of participants with an AA genotype of rs13294, AA and AG genotypes were grouped as AX. In both Figs. 7(a) and (b), the participants from the low sensitivity group have higher insula and amygdala GMD values than those in the high sensitivity group (p-value = 0.0014 for insula, p-value = 0.0037 for amygdala). The main effects of genotype show that the GMD of the insula and amygdala of the AX group (dark yellow) is lower than the GG group (light yellow) with p-values = 0.0137 and 0.0474 respectively. Moreover, a significant interaction effect can be found between the two independent variables with p-values = 0.0463 and 0.0476 for the insula and amygdala respectively.

**Fig. 7.**
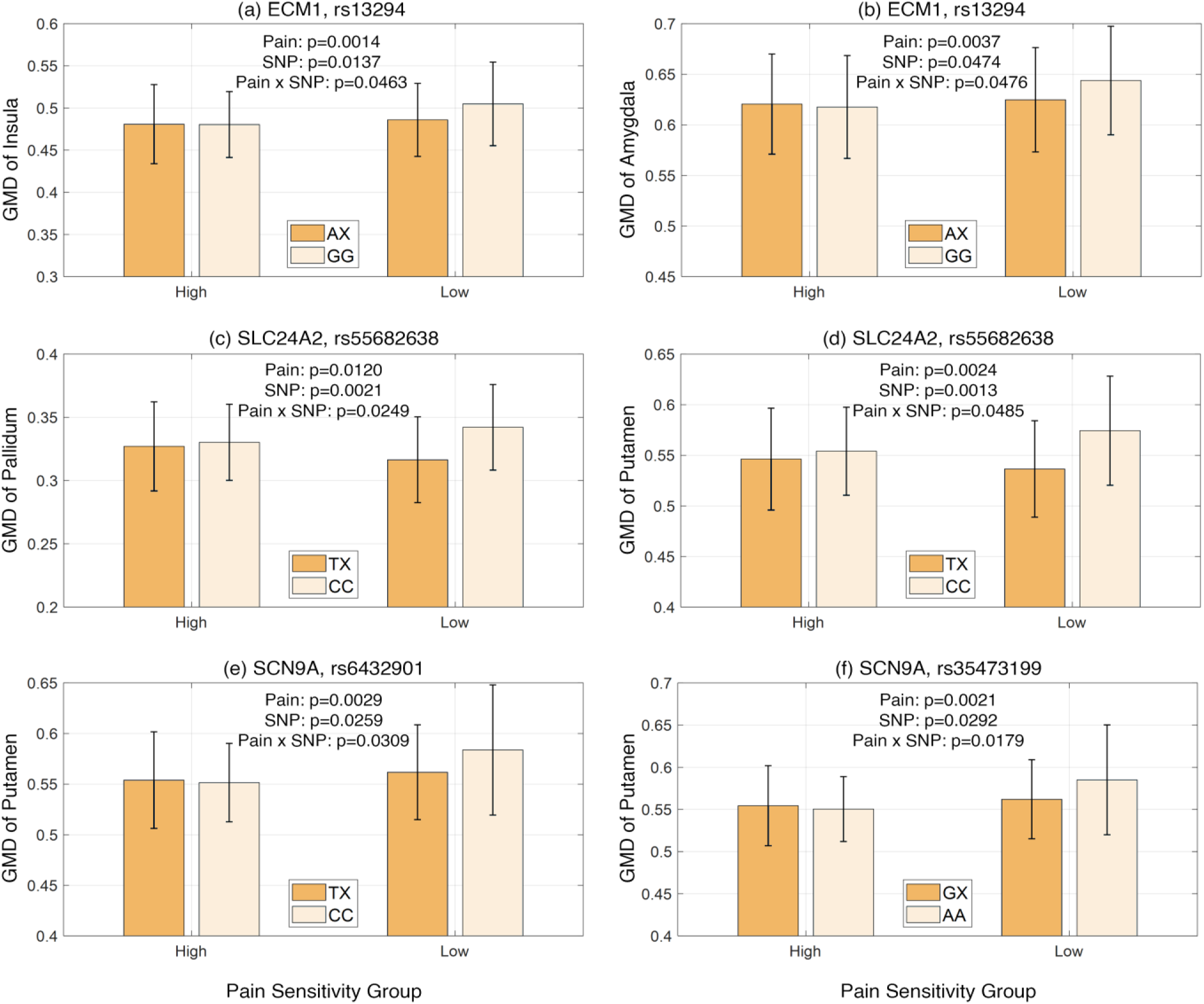
Statistical analysis of identified pain sensitivity-related brain region GMD about corresponding SNPs and pain sensitivity groups. Two-way ANOVA was conducted to examine the effect of each SNP and pain sensitivity on the GMD of these regions. The p-values for the main effect and interaction effect are shown in the plot. The bars and error bars indicate the mean and standard deviation of GMD.

For the *SLC24A2* gene, in total 51 SNPs can be found in our genotyping dataset. We examined the effect of these target SNPs and pain sensitivity groups on the GMD from ROIs with significant differences. One SNP rs55682638 demonstrated significant results as shown in Figs. 7(c) and (d). For both results from Figs. 7(c) and (d), TT and TC genotypes are grouped with in total of 50 participants. The CC group has 315 participants. From the results, we can see that there is a significant main effect of pain sensitivity on GMD of pallidum and putamen respectively (p-value = 0.0120 and 0.0024). This indicates that participants of the high pain sensitivity group have lower GMD of both pallidum and putamen. Similarly, genotype also shows a significant main effect on the GMD value, with p-values of 0.0021 and 0.0013 for rs55682638 on the GMD of pallidum and putamen. Moreover, the interaction between pain sensitivity and genotype is significant, with p-values of 0.0249 and 0.0485 respectively for the two ROIs. The significant interaction suggests that the influence of genotype on GMD depends on pain sensitivity and vice versa.

Similarly, after performing the two-way ANOVA of 27 SNPs from *SCN9A* on GMD of pain matrix ROIs, two SNPs rs6432901 and rs35473199 show significant results between pain sensitivity and genotyping groups. The results are shown in Figs. 7(e) and (f). Both significant results are related to the GMD of putamen. Firstly, Figs. 7(e) and (f) indicate that participants with high pain sensitivity have significantly lower putamen GMD values for both target SNPs (p-value = 0.0029 and 0.0021 for rs6432901 and rs35473199 respectively). In Fig. 7(e), TT and TC genotypes were grouped as TX, similar to the previous analysis. The GMD of putamen of the TX group (dark yellow, n = 236) was significantly lower than the CC group (light yellow, n = 124) of rs6432901 (p-value = 0.0259). In Fig. 7(f), we can also find that the putamen GMD of the GX group (dark yellow, n = 242) was significantly lower than the AA group (light yellow, n = 120) of rs35473199 (p-value = 0.0292). Moreover, a significant interaction effect can be found for both SNPs with p-values = 0.0309 and 0.0179 respectively.

### 4.7. Mediation effects of GMD on the relationship between SNP and pain threshold

After conducting the two-way ANOVA analyses, we identified significant interaction effects among the genotypes of critical SNP, GMD, and pain sensitivity. Subsequently, through mediation analysis, we discovered a significant indirect effect of the SNP genotype on pain sensitivity measurements (pain threshold) mediated by GMD. This effect was observed in 5 SNP-ROI pairs (rs13294-insula, rs6432901-putamen, rs35473199-putamen, rs55682638-pallidum, rs55682638-putamen). The mediation model results are illustrated in Fig. 8.

**Fig. 8.**
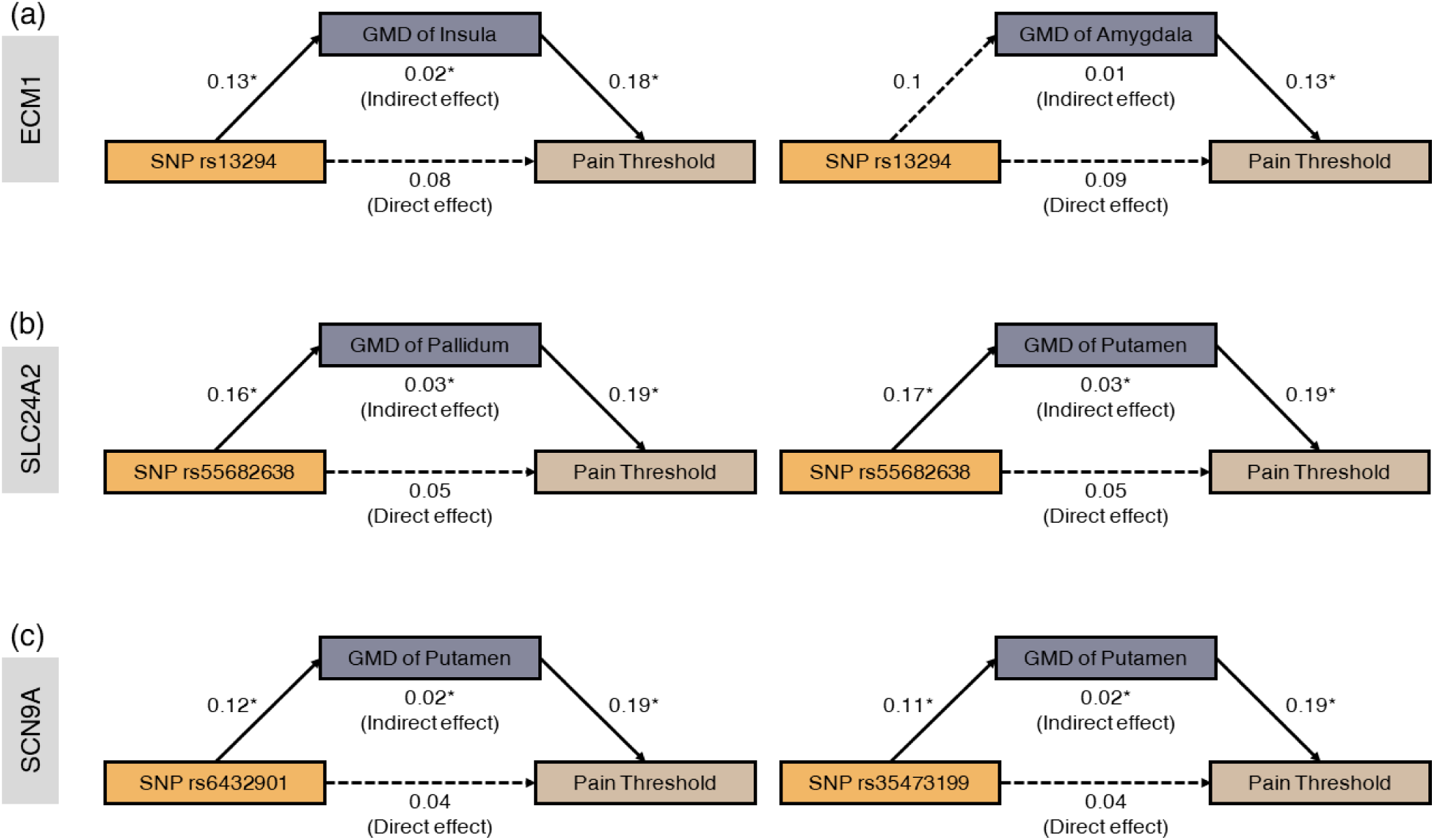
Mediation analysis results for three genes *ECM1*, *SLC24A2* and *SCN9A*. The mediation analysis reveals that the relationship between SNP genotype and pain sensitivity (pain threshold) was indirected-only mediated by GMD of corresponding brain regions.

We observed significant associations among the SNP genotype of the *ECM1* gene, pain threshold, and the GMD of the left insula. The mediation analysis demonstrated that the SNP rs13294 had an indirect effect (β = 0.02, SE = 0.05, CI = [0.009, 0.111], p = 0.013) but not a direct effect (β = 0.08, SE = 0.07, CI = [-0.023, 0.18], p = 0.12) on pain threshold through the GMD of the insula (Fig. 8(a) left).

Moreover, concerning the *SLC24A2* gene, the mediation analysis indicated that the SNP rs55682638 had an indirect effect (β = 0.03, SE = 0.01, CI = [0.027, 0.186], p = 0.003) but not a direct effect (β = 0.05, SE = 0.05, CI = [-0.05, 0.143], p = 0.087) on pain threshold mediated by the GMD of pallidum (Fig. 8(b) left) and similarly for the same SNP through the GMD of putamen (indirect effect: β = 0.03, SE = 0.02, CI = [0.027, 0.191], p = 0.003; direct effect: β = 0.05, SE = 0.04, CI = [-0.047, 0.145], p = 0.057) (Fig. 8(b) right).

Regarding the *SCN9A* gene, the mediation analysis revealed that the SNP rs6432901 also had an indirect effect (β = 0.02, SE = 0.03, CI = [0.006, 0.116], p = 0.027) but not a direct effect (β = 0.04, SE = 0.04, CI = [-0.064, 0.139], p = 0.077) on pain threshold through the GMD of the putamen (Fig. 8(c) left). Similar findings were observed for the SNP rs35473199 (indirect effect: β = 0.02, SE = 0.02, CI = [0.005, 0.125], p = 0.023; direct effect: β = 0.04, SE = 0.06, CI = [-0.067, 0.136], p = 0.082) (Fig. 8(c) right). These findings demonstrate that the relationships between SNP genotype and pain sensitivity were partially mediated by GMD.

## 5. Discussion

This study investigates the genetic basis of pain sensitivity individual differences using a combination of genomics, transcriptomics, and brain neuroimaging. We first identified 15 brain regions with significant GMD differences between high and low pain sensitivity groups (Fig. 2). Next, PLS analysis found positive correlations between gene expression patterns and GMD differences across six donors in the left hemisphere (Fig. 3). Clustering and enrichment analysis of 27 key differentially expressed genes revealed distinct patterns and functional associated with GMD differences (Figs. 4 and 5). We further examined the expression levels of these genes across various brain regions using a pain-related gene set (Fig. 6). Additionally, significant interactions were observed between SNP genotypes from *ECM1*, *SLC24A2*, and *SCN9A* and pain sensitivity on GMD differences (Fig. 7). Mediation analysis demonstrated that these SNP and pain threshold were mediated by GMD (Fig. 8). In conclusion, we propose a interpretation model that integrates genomic, transcriptomic, and imaging features to describe how these factors shape individual differences in pain sensitivity (Fig. 9).

**Fig. 9.**
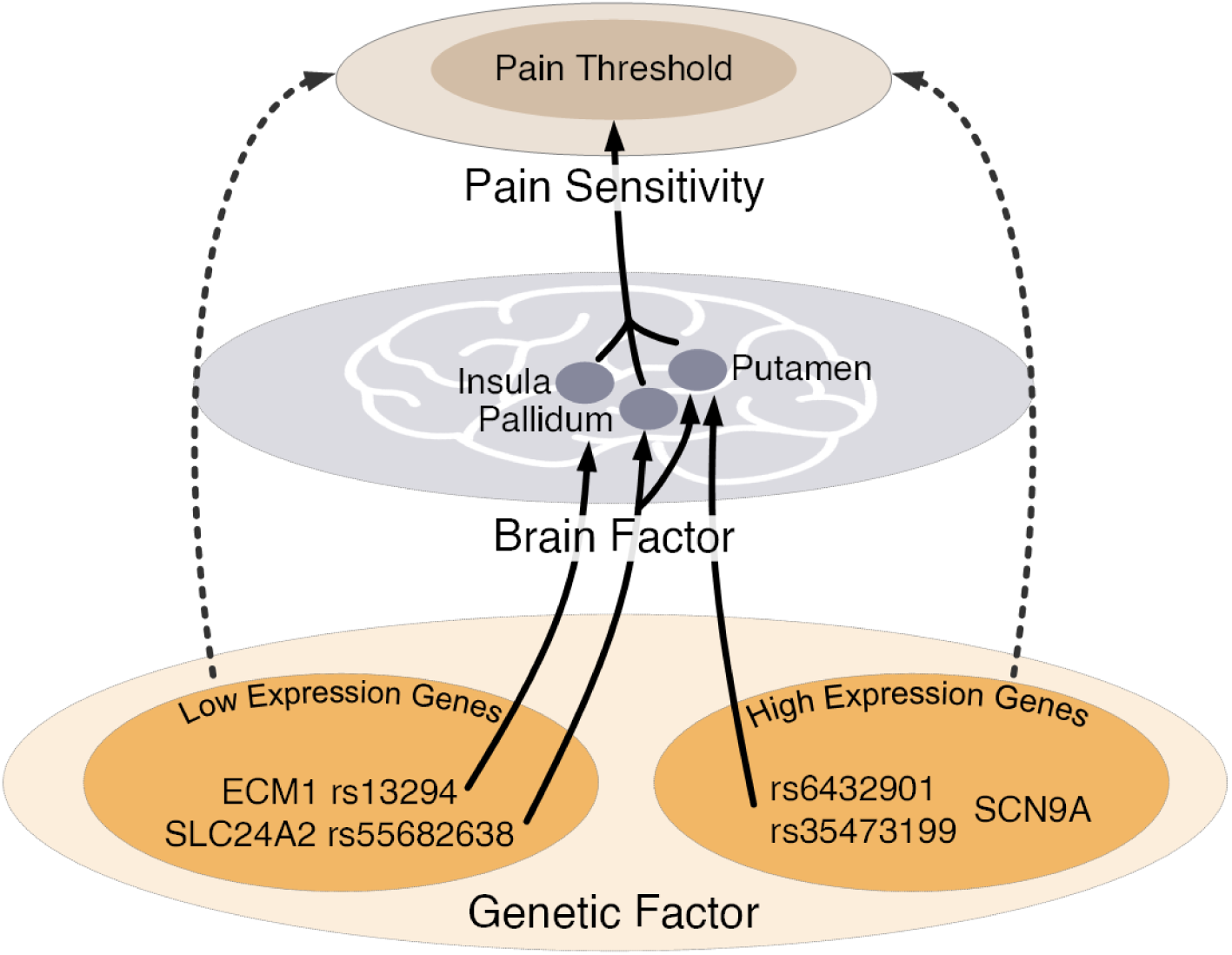
Interpretation model to explain the mechanism underlying the interplay of genetic factors, brain factors and pain sensitivity. From the genetic factor perspective, there exist two gene groups with different expression patterns. Meanwhile, the genotypes of these genes influence the pain sensitivity in an indirect-only way via GMD of brain regions from basal ganglia.

### 5.1. Brain structural phenotype

Pain processing is associated with multiple brain regions.^14^ In Fig. 2, in total 15 brain regions have been identified to be significantly associated with pain sensitivity, such as amygdala, basal ganglia, insula, and thalamus. There are multiple identified regions belonging to traditional pain matrix or pain-related regions.^5,11,63^ Previous work highlighted the amygdala’s role in connecting pain perception with emotional experience.^63^ Evidence has been shown that the basal ganglia structures is activated during pain experiences, as well as their involvement in modulating pain perception.^11^ These regions collaborate to receive, process and integrate pain signals, contributing to the subjective experience of pain.

### 5.2. Key genes enrichment analysis

After using PLS regression analysis and validation as shown in Fig. 3, we identified 27 key genes associated with the GMD differences between high and low pain sensitivity groups. As shown in Fig. 4 (a), the gene clustering illustrates that these genes can notably be divided into two main groups. We also observed that within each group, the correlations between genes are mostly positive while between groups correlations are mostly negative.

Fig. 5(b) shows that group 1 genes are significantly enriched in cellular components related to the main axon, VGSC and the node of Ranvier. These findings are biologically reasonable, as pain perception is predominantly mediated by specialized nerve fibers termed nociceptors. The main axons carry sensory information from the peripheral to the central nervous system, signaling potential skin-damaging stimuli. VGSCs serve as primary targets for analgesic drugs, emphasizing their importance in pain modulation.^10^ Mutations in VGSC genes, like NaV1.7, have been linked to altered pain sensitivity. Global Nav1.7-knockout mice exhibited a phenotype resembling human congenital indifference to pain, supporting the role of these genes in pain perception.^4,7^

Regarding molecular function, the high expression genes were also enriched in functions related to monovalent inorganic cation transmembrane transport, VGSC activity, cation channel activity, and sodium/metal ion transmembrane transport activities. The above enrichment functions are relevant to the activation and transport of ion channels, which play important roles in neuronal excitability and neurotransmission, including processes involved in pain perception. For example, transient receptor potential (TRP) channels, activated by stimuli like heat and protons, can significantly impact pain sensitivity when their activity or expression is altered.^6,46^ In contrast, two of the genes (*SCN9A* and *S100A10*) showed significant differences in gene expression patterns, but no significant enrichment in molecular functions. These findings indicate that these clustered gene groups might correspond to functional disparities linked to pain sensitivity, due to differences in their activities and processes.

Further analysis integrated these 27 genes with the pain-related gene database, identifying 8 key genes. Fig. 6 shows the gene expression level of these 8 genes across different brain regions. A polarized expression pattern can be observed in basal ganglia regions, including the amygdala, caudate, putamen, pallidum, and thalamus. The basal ganglia are known to be involved in pain perception and modulation.^5,9,27,59^ The results further suggest that the distinct transcriptional alternations may contribute to variations observed in the imaging structural features between different pain sensitivity groups.

### 5.3. SNP statistical analysis

The statistical analysis shown in Figs. 7(a) and (b) demonstrate the association between the *ECM1* gene SNP rs13294 with the GMD of the amygdala and insula, respectively. From both figures, the GMD between the AX and GG genotypes varies significantly between individuals with high and low pain sensitivity groups. Specifically, individuals with the GG homozygotes of rs13294 exhibit lower amygdala/insula GMD in the high pain sensitivity group and higher GMD in the low pain sensitivity group. These findings are promising, aligning with previous studies on pain perception and modulation.^53,56,63^ For instance, Lu and colleagues^36^ studied the anterior insular cortex’s role in integrating various information sources to create pain awareness. ^36^ Similarly, other research has shown the amygdala’s function as a central hub integrating sensory input from nociceptive pathways with emotional and cognitive processes.^53,63^ From genetic aspects, studies have linked the rs13294 variant in the *ECM1* gene to chronic musculoskeletal pain.^61^ Laminin β1 (LAMB1) is a key element of the extracellular matrix protein. Previous study has shown that LAMB1 in the ACC was significantly downregulated upon peripheral neuropathy and knocking down of LAMB1 in the ACC can modify mice pain sensitivity, suggesting its role as a modulator in pain signaling pathways.^32^

Figs. 7(c) and (d) show that the interaction between genotype and pain sensitivity significantly influences GMD in the pallidum and putamen. Individuals with the CC homozygotes of rs55682638 exhibit higher GMD levels in both high and low pain sensitivity groups. Research on the *SLC24A2* gene indicates its potential role in pain modulation and neurotransmitter regulation.^58,67^ For instance, Zhou et al. demonstrated that increased *SLC24A2* expression could decrease thermal pain sensitivity and the expression of inflammatory factors in rats with chronic constriction injury.^58^ Additionally, the basal ganglia, including the pallidum and putamen, are involved in pain processing pathways.^5,9^ The findings establish a clear link between the SNP rs55682638, the pallidum and putamen concerning pain sensitivity, suggesting that *SLC24A2* and the basal ganglia might offer potential targets for pain modulation strategies.

The results from Figs. 7(e) and (f) indicate significant correlations between two *SCN9A* gene SNPs (rs6432901 and rs35473199) and the GMD of putamen.^20,51^ The two-way ANOVA analysis demonstrates an interaction effect between genotype and pain sensitivity concerning putamen GMD. Specifically, individuals with the CC genotype for both SNPs show lower putamen GMD in the high pain sensitivity group and higher GMD in the low pain sensitivity group. *SCN9A* encodes the Nav1.7 sodium channel important in pain perception, and the putamen is part of the pain matrix, containing opioid receptors and neurons responsive to various pain stimuli.^56^

### 5.4. Interpretation model

In Fig. 8, the mediation analysis revealed possible interactions among genotype, brain GMD features and individual pain sensitivity. The analysis suggests that certain SNP genotypes regulate pain sensitivity (pain threshold) by modulating the multiple brain structural features. To summarize our results, we propose an interpretation model in that integrates the roles of contribution from genetic and brain factors in influencing pain sensitivity (Fig. 9). Individual pain sensitivity is affected by both genetic and brain factors, particularly in regions like the basal ganglia, insula, and other traditional pain matrix areas. Two groups of genes were identified with polarized expression patterns in the basal ganglia that are related to GMD and pain sensitivity. *ECM1* gene and *SLC24A2* gene are low expressed in the basal ganglia, which mainly regulate the insula, pallidum, and putamen to influence pain sensitivity. On the contrary, the *SCN9A* gene is highly expressed in the basal ganglia, which primarily contributes to putamen GMD to regulate pain sensitivity.

The statistical and mediation analyses indicate that genotype regulates pain sensitivity by modulating the intrinsic brain structural GMD features in these regions. This suggests two distinct ways in which genetic patterns can influence pain sensitivity through their effects on basal ganglia structure and function. Our proposed model provides a deeper understanding on the interplay between genetics, brain structure, and pain sensitivity. Further research is needed to elucidate the underlying mechanisms behind these two genetic pathways of pain regulation.

### 5.5. Limitation

Several limitations should be considered when interpreting the results of this study. First, the sample size was relatively small, and future studies with larger sample sizes and external validation are needed to further verify GMD differences across individuals. Second, the gene expression data and neuroimaging data were collected from different participants. Though the genotype SNP and the brain neuroimaging data are collected from the same population, the transcriptional gene expression data are not consistent. The gene expression profiles in AHBA are influenced by factors such as age, sex, and ethnicity, which may introduce bias and limit the generalizability of the findings to diverse populations. Therefore, multi-omics datasets from the same subjects are more promising to provide reliable results. Third, our mediation analysis model only shed light on potential causality. However, to determine the causal relationship, further validation through animal experiments is required.

## 6. Conclusion

In conclusion, we collected pain threshold, brain MRI data and genotype SNP data from 432 subjects. We first used MRI data to analyze the brain GMD features that related to pain sensitivity. Next, we used PLS regression by integrating the AHBA transcriptional gene expression dataset and GMD features difference between high and low pain sensitivity groups to locate key genes that may regulate related brain regions. Enrichment and clustering analysis were used to reveal the molecular functions and cellular components of two key gene sets with different expression patterns. Moreover, a series of statistical analyses using genotype SNP data demonstrated potential key loci that may related to brain structural GMD features and pain sensitivity. Mediation analysis revealed the contribution of genetic factors and brain factors to pain sensitivity. In the end, we proposed an interpretation model to summarize our results. Our study provides valuable insights into the transcriptomic and genomic signature of GMD patterns that may shape individual pain sensitivity.

## Acknowledgments

This work was supported in part by the National Natural Science Foundation of China (62201356, 32361143787), in part by the Guangdong Basic and Applied Basic Research Foundation (2021A1515110694), in part by the Shenzhen-Hong Kong Institute of Brain Science-Shenzhen Fundamental Research Institutions (2022SHIBS0003), and in part by Medicine Plus Program of Shenzhen University (2024YG021).

