## Supplementary Material for "Genomic and transcriptomic profiles influence on brain morphology and their interactions with pain sensitivity individual differences" (from another version of this preprint: version 1, 2024-07-30)

### 1. Gene set from PLS and clustering.

We identified genes that significantly contributed to PLS1 consistently present across all six donors using the bootstrapping method. More specifically, we identified a gene set that is associated with pain sensitivity-related GMD alternation for each donor. In order to obtain a relatively stable result, we only selected genes are were identified from all six donors for further analysis. In the end, 27 key genes were identified to be significantly associated with differences in GMD between two pain sensitivity groups and were used. Fig. S1 shows the Venn diagram of the identified gene numbers and their overlapping from all six donors. Next, we used a k-means clustering method on these 27 key genes. The 27 significant genes were divided into two groups: group 1 with 21 genes, and group 2 with 6 genes. The gene names, Ensembl ID, basic gene description, chromosome ID, gene type and their group ID after the clustering are shown in Table S1.

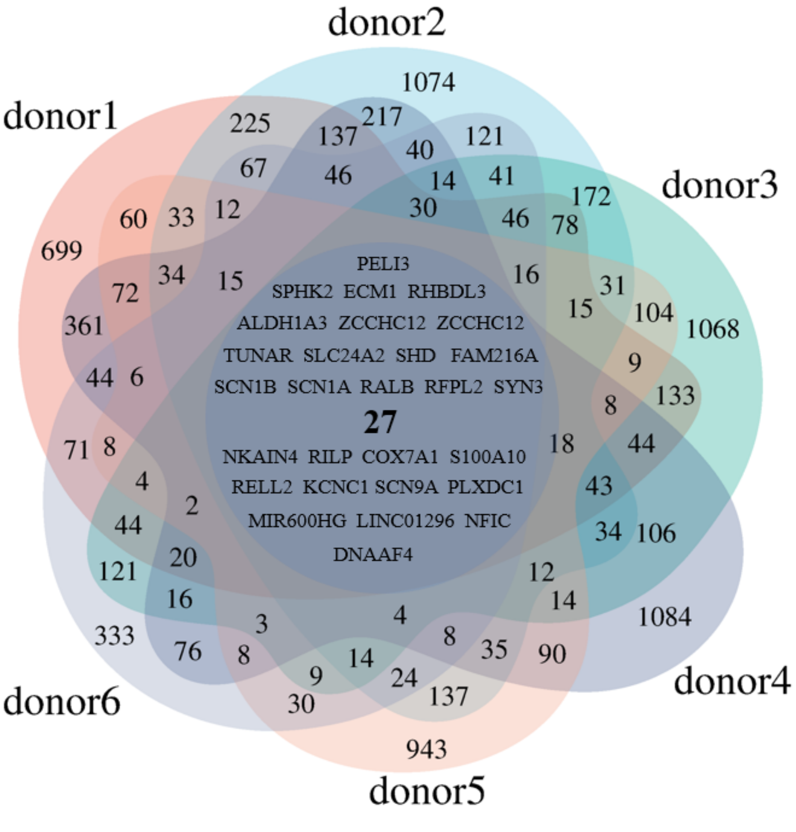


Fig. S1. The Venn diagram of the identified genes from all six donors.

Table S1. The gene names, Ensembl ID, basic gene description, chromosome ID, gene type and their group ID after the clustering.

| Gene name | Ensembl ID | Gene description | Chr | Gene type | Group |
|---|---|---|---|---|---|
| ALDH1A3 | ENSG00000184254 | Aldehyde Dehydrogenase 1 Family Member A3 | 15 | Protein Coding | 1 |
| COX7A1 | ENSG00000161281 | Cytochrome C Oxidase Subunit 7A1 | 19 | Protein Coding | 1 |
| DNAAF4 | ENSG00000256061 | Dynein Axonemal Assembly Factor 4 | 15 | Protein Coding | 2 |
| ECM1 | ENSG00000143369 | Extracellular Matrix Protein 1 | 1 | Protein Coding | 1 |
| FAM216A | ENSG00000204856 | Family With Sequence Similarity 216 Member A | 12 | Protein Coding | 1 |
| KCNC1 | ENSG00000129159 | Potassium Voltage-Gated Channel Subfamily C Member 1 | 11 | Protein Coding | 1 |
| LINC01296 | ENSG00000225210 | Double Homeobox A Pseudogene 9 | 14 | Pseudogene | 1 |
| MIR600HG | ENSG00000236901 | MIR600 Host Gene | 9 | RNA Gene | 1 |
| NFIC | ENSG00000141905 | Nuclear Factor I C | 19 | Protein Coding | 1 |
| NKAIN4 | ENSG00000101198 | Sodium/Potassium Transporting ATPase Interacting 4 | 20 | Protein Coding | 2 |
| PELI3 | ENSG00000174516 | Pellino E3 Ubiquitin Protein Ligase Family Member 3 | 11 | Protein Coding | 1 |
| PLXDC1 | ENSG00000161381 | Plexin Domain Containing 1 | 17 | Protein Coding | 1 |
| RALB | ENSG00000144118 | RAS Like Proto-Oncogene B | 2 | Protein Coding | 1 |
| RELL2 | ENSG00000164620 | RELT Like 2 | 5 | Protein Coding | 1 |
| RFPL2 | ENSG00000128253 | Ret Finger Protein Like 2 | 22 | Protein Coding | 1 |
| RHBDL3 | ENSG00000141314 | Rhomboid Like 3 | 17 | Protein Coding | 1 |
| RILP | ENSG00000167705 | Rab Interacting Lysosomal Protein | 17 | Protein Coding | 1 |
| S100A10 | ENSG00000197747 | S100 Calcium Binding Protein A10 | 1 | Protein Coding | 2 |
| SCN1A | ENSG00000144285 | Sodium Voltage-Gated Channel Alpha Subunit 1 | 2 | Protein Coding | 1 |

| | | | | | |
|---|---|---|---|---|---|
| SCN1B | ENSG00000105711 | Sodium Voltage-Gated Channel Beta Subunit 1 | 19 | Protein Coding | 1 |
| SCN9A | ENSG00000169432 | Sodium Voltage-Gated Channel Alpha Subunit 9 | 2 | Protein Coding | 2 |
| SHD | ENSG00000105251 | Src Homology 2 Domain Containing Transforming Protein D | 19 | Protein Coding | 1 |
| SLC24A2 | ENSG00000155886 | Solute Carrier Family 24 Member 2 | 9 | Protein Coding | 1 |
| SPHK2 | ENSG00000063176 | Sphingosine Kinase 2 | 19 | Protein Coding | 1 |
| SYN3 | ENSG00000185666 | Synapsin III | 22 | Protein Coding | 1 |
| TUNAR | ENSG00000250366 | TCL1 Upstream Neural Differentiation-Associated RNA | 14 | Protein Coding | 2 |
| ZCCHC12 | ENSG00000174460 | Zinc Finger CCHC-Type Containing 12 | X | Protein Coding | 2 |

### 2. Pain-related gene identification

As presented earlier, we have identified 27 key genes from our imaging transcriptomics analysis. By taking the intersection set of these 27 key genes and the aforementioned pain-related genes set from multiple database sources (including the Human Pain Genetics Database45, Pain Genes Database46, Gene Ontology search employing the keyword “pain”, and Pain Research from https://www.iasp-pain.org/publications/pain-research-forum/), we further narrow our analysis down to 8 significant pain-related genes: ECM1, SCN1A, SCN1B, SLC24A2, SPHK2, SYN3, S100A10 and SCN9A. Table S2 shows the genes intersection of these 27 key genes and four pain-related databases. Fig. S2 shows the Venn diagram of the intersection.

Table S2. Genes intersection of these 27 key genes and four pain-related databases.

| **Database** | **Intersect with 27 key genes** | **Identified gene number / All gene number in this database** |
|---|---|---|
| Human Pain Genetics Database | ECM1, SCN1A, SCN9A, SYN3 | 4/480 |
| Pain Genes Database | S100A10, SCN9A, SPHK2 | 3/421 |
| Gene Ontology “pain” | SCN1A, SCN1B, SCN9A, SLC24A2 | 4/120 |
| Pain Research Forum | SCN1A, SCN9A | 2/94 |

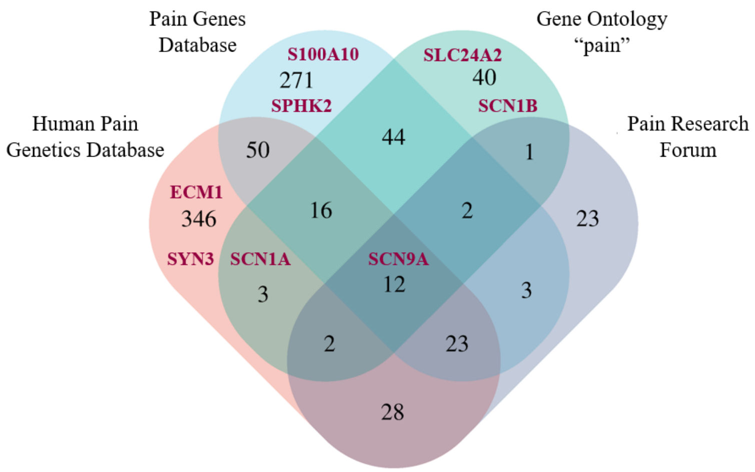


Fig. S2. The Venn diagram of the intersection of the 27 key genes and 4 pain-related databases.

### 3. Clustering results from 8 key gene

We used a *k*-means clustering method on the 8 key genes (ECM1, SCN1A, SCN1B, SLC24A2, SPHK2, SYN3, S100A10 and SCN9A) based on their expression value on the whole-brain 45 ROIs and 10 pain matrix ROIs respectively. The clustering results are shown in Fig. S3. To compare the clustering effect with different k, the cluster numbers were chosen as 2 and 3 and the silhouette coefficients are also shown respectively.

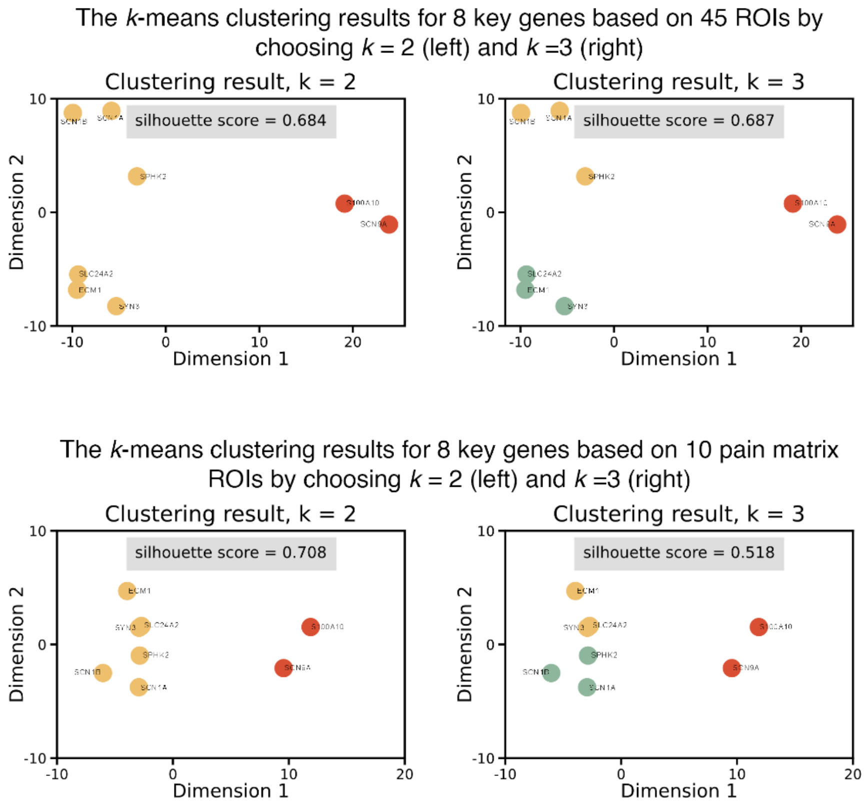


Fig. S3. The clustering results based on 8 identified key genes. The upper two sub-figures are based on whole-brain 45 ROIs and the lower two sub-figures are based on pain matrix ROIs.
